# Start from Scratch: Precisely Identify Massive Active Enhancers by Sequencing

**DOI:** 10.1101/604686

**Authors:** Xiusheng Zhu, Lei Huang, Qing Li, Yubo Zhang

## Abstract

Enhancer loci identified by ChIP-Seq or other experimental methods occupy hundreds of base pairs on the genome. It is the paradox comparing with the motif analysis, which usually contains only a few or tens nucleotides, achieved by bioinformatics analysis. To address this issue, we develop an experimental method, termed as massive active enhancer sequencing (MAE-Seq), to designate active enhancer sequences from arbitrary sources of 25bp random DNA libraries. These sequences are constructed in a mini-promoter vector with fluorescent reporter. After transfection, positive cells are sorted out and for sequencing. With the results, we successfully identify hundreds known accurate active enhancer sequences as expected. Besides that, large amounts of unmarked regulatory elements (UREs) without epigenetic features are also been spotted. In conclusion, MAE-Seq would be useful to refine enhancer sequences and precisely annotate the genome in eukaryotes.

## Introduction

Enhancers are critical regulatory elements determining precisely spatiotemporal gene expression^1,2^. Characterizing the genomic feature of this kind of elements is one of the major tasks for ENCODE (Encyclopedia of DNA elements) project^1,2^. Different methods have been developed for the purpose, such as ChIP-seq (chromatin immunoprecipitation)^3,4^, DNase-seq (DNase I hypersensitive sites sequencing)^5^, FAIRE-seq (Formaldehyde-Assisted Isolation of Regulatory Elements)^6^, ATAC-seq (Assay for Transposase-Accessible Chromatin using sequencing)^7^ and STARR-seq (self-transcribing active regulatory region sequencing)^8,9^. Collectively, these methods rely on chromatin feature of histone modification^10^ or chromatin accessibility^11^, which are considered as hallmarks to characterize enhancer loci across genome. However, due to the chromatin characteristics and analysis strategies, the annotation of such elements in ENCODE is not accurate enough^1,2,12^.

Usually, defined regulatory elements span hundreds (or even thousands) of base pairs (bp) on the genome^13^ (Fig. 1A). For example, the coverage regions of H3K27ac ChIP-seq datasets are ranging from 200 to 6×10^4^ bp^10^ (such as newly defined super-enhancers^14,15^). These regions might contain multiple signals^14^. At the same time, with motif analysis on these different genomic datasets, the core regions turn out only span a few or tens bp at each locus^16^. It is well known that the nucleosome consisted by nine subunits of histones and one piece of 147 bp DNAs. Therefore, it might be an impossible mission to precisely define core enhancer sequences across the genome with available technologies experimentally.

**Figure 1.**
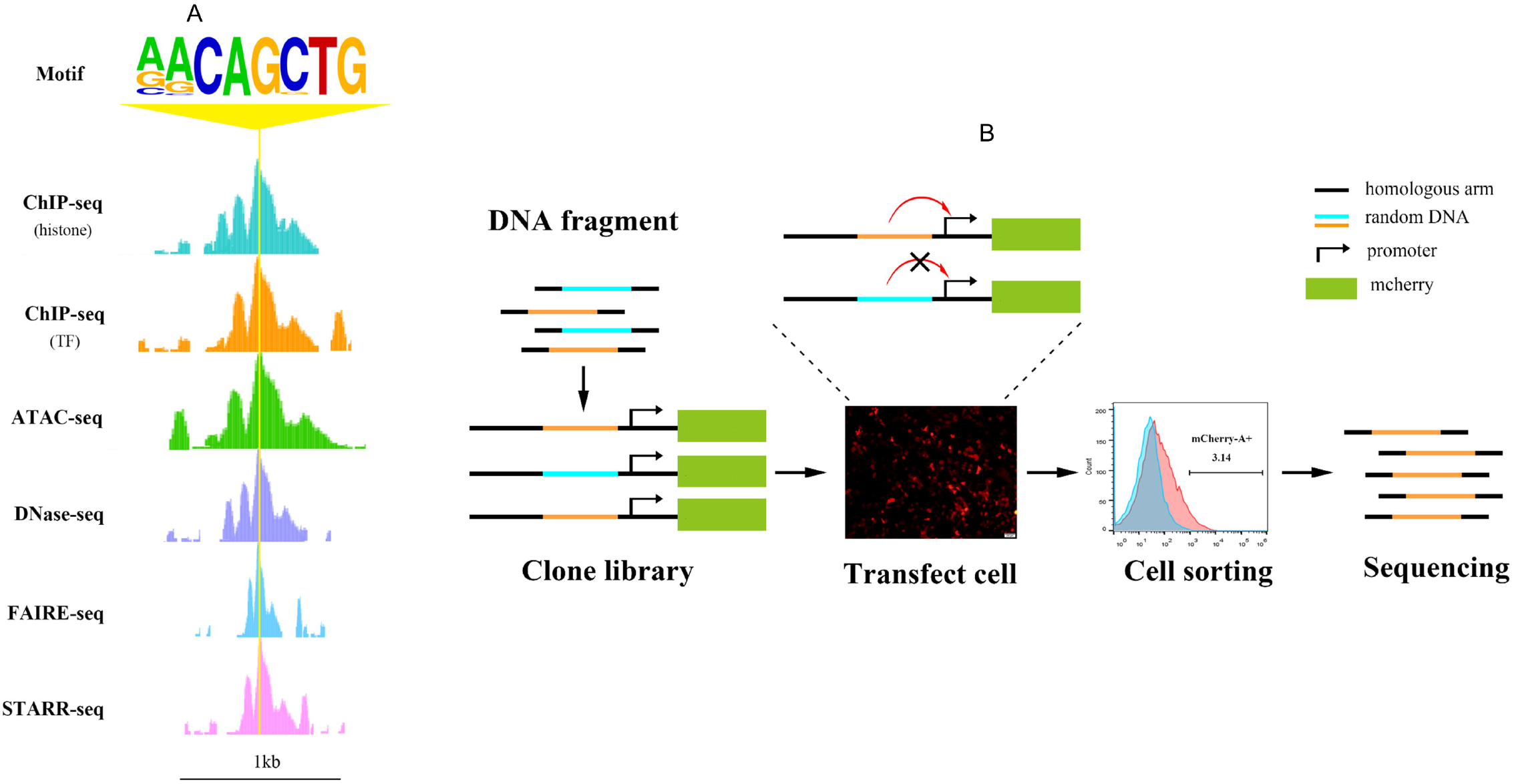
Overview of the MAE-seq method. A: Regulatory elements defined by existing technologies. B: MAE-Seq workflow. Random DNAs with homologous arm are constructed in a mini-promoter vector with fluorescent reporter. After transfection, cells are sorted out and for HTS.

In the study, we develop a new technology termed as MAE-Seq (Massive Active enhancers by sequencing) to identify the precise position of enhancers. Random DNA fragments are introduced into a mini-promoter vector with fluorescent reporter. After transfection, positive cells are sorted out by flow cytometry. We test a series of fragments in different lengths (20-50bp) and finally determine 25bp random DNA library for enhancers identification. Base on MAE-seq, thousands of core enhancers with length of 25bp are spotted across genome.

## Results

### MAE-Seq development

In this study, we develop MAE-Seq which relies on random DNA libraries and fluorescence-assisted screening (Fig. 1B). In brief, random DNA fragments are synthesized, and constructed in a mini-promoter vector with fluorescent reporter. Then the clone library is transfected into cultured cells. Positive cells are sorted out by flow cytometry and for Illumina high-throughput sequencing (HTS).

To figure out proper length of random DNA fragments for achieving high unique mapping rate, fragments with different length (20~60bp) are generated from human reference genome and aligned. Results show that, the length of fragments does have a significant effect on the unique mapping rate. The rate is less than 20% when it is 20 bp, while the rate of 25 bp elevates to more than 60% and reached a plateau stage of about 70% between 30 bp and 40 bp (Fig. 2A). Meanwhile, in order to test whether it affects the fluorescence signal of the reporting vector system, the fluorescence intensity (FI) of the positive cells with different length of fragments is analyzed. Results reveal that, the length of small fragments within 50 bp does not affect the sensitivity of the fluorescence reporting system (Fig. 2B). Though the FI of cells with 25 bp fragments is slightly high, there is no significant difference in FI compared with cells with fragments which length is shorter than 50 bp. Therefore, 25 bp fragments are selected to construct random DNA libraries and screen active enhancers (Fig. 2C and 2D).

**Figure 2.**
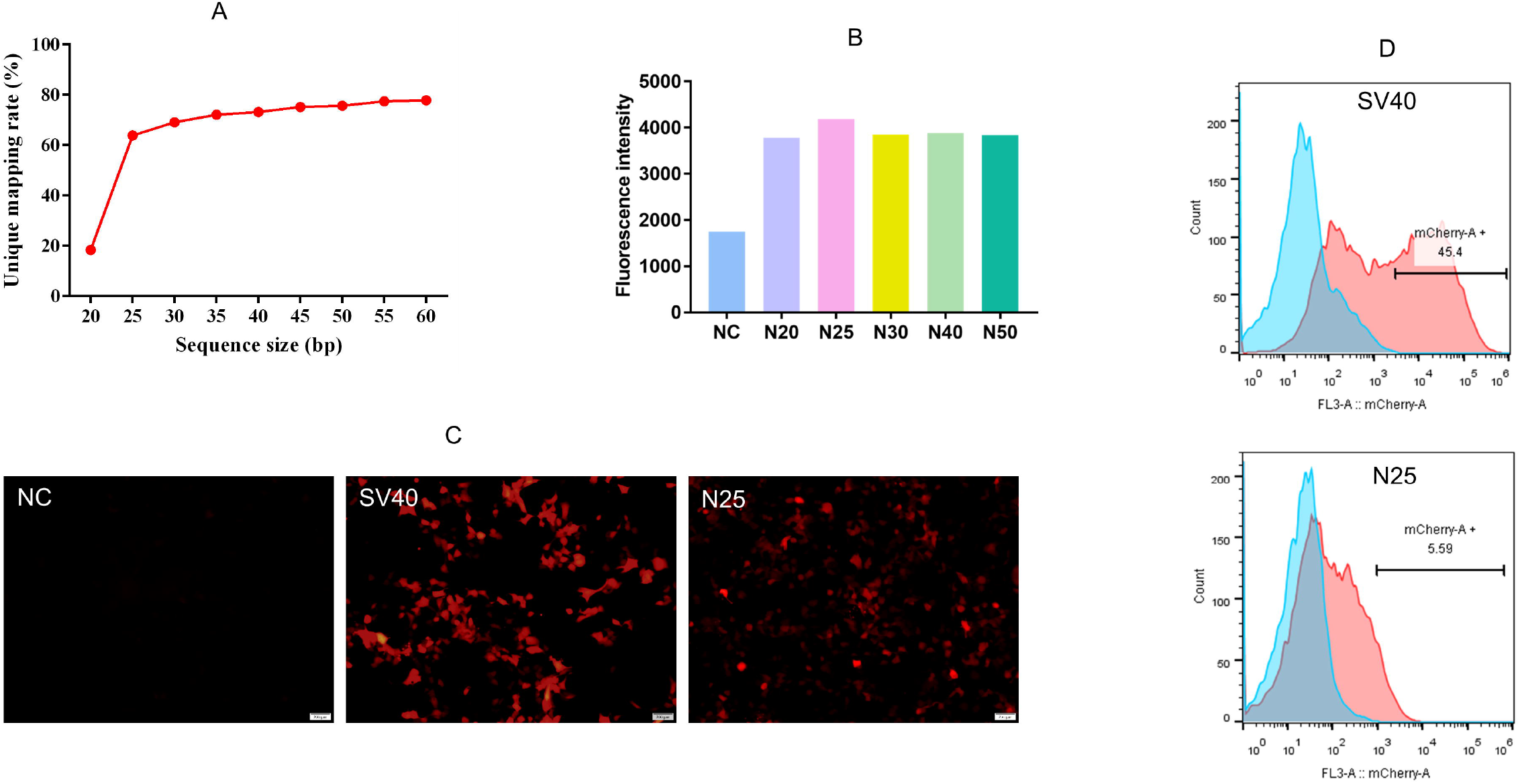
Length test and cell sorting. A: Simulation by intercepting random fragments with different lengths versus unique mapping rate. B: Intensity analysis of the positive cells. C: Fluorescent signals. NC: empty vector, SV40: positive control, N25: 25bp random DNA library. D: Statistical peaks. Light blue: negative cell population, Calibrated orange: positive cell population.

### Identified active enhancers

After quality control and removing duplication (Fig. 3A), 0.5 million valid reads are achieved (table 1). They are aligned to the reference genome (hg19), 9771 loci which corresponded to 12412 unique mapping reads are obtained (table 2). Among them, 49.7% and 45.1% loci distribute to intergenic and intron regions, respectively (Fig. 3B) (Table S1). Moreover, 39.23% or 33.70% of them are located in the range of 5-50 kb or 50-200kb upstream / downstream of TSS locus (Fig. 3C), which are the high frequency regions where conventional or distal enhancers (CEnh or DEnh) appeared^17^.

**Figure 3.**
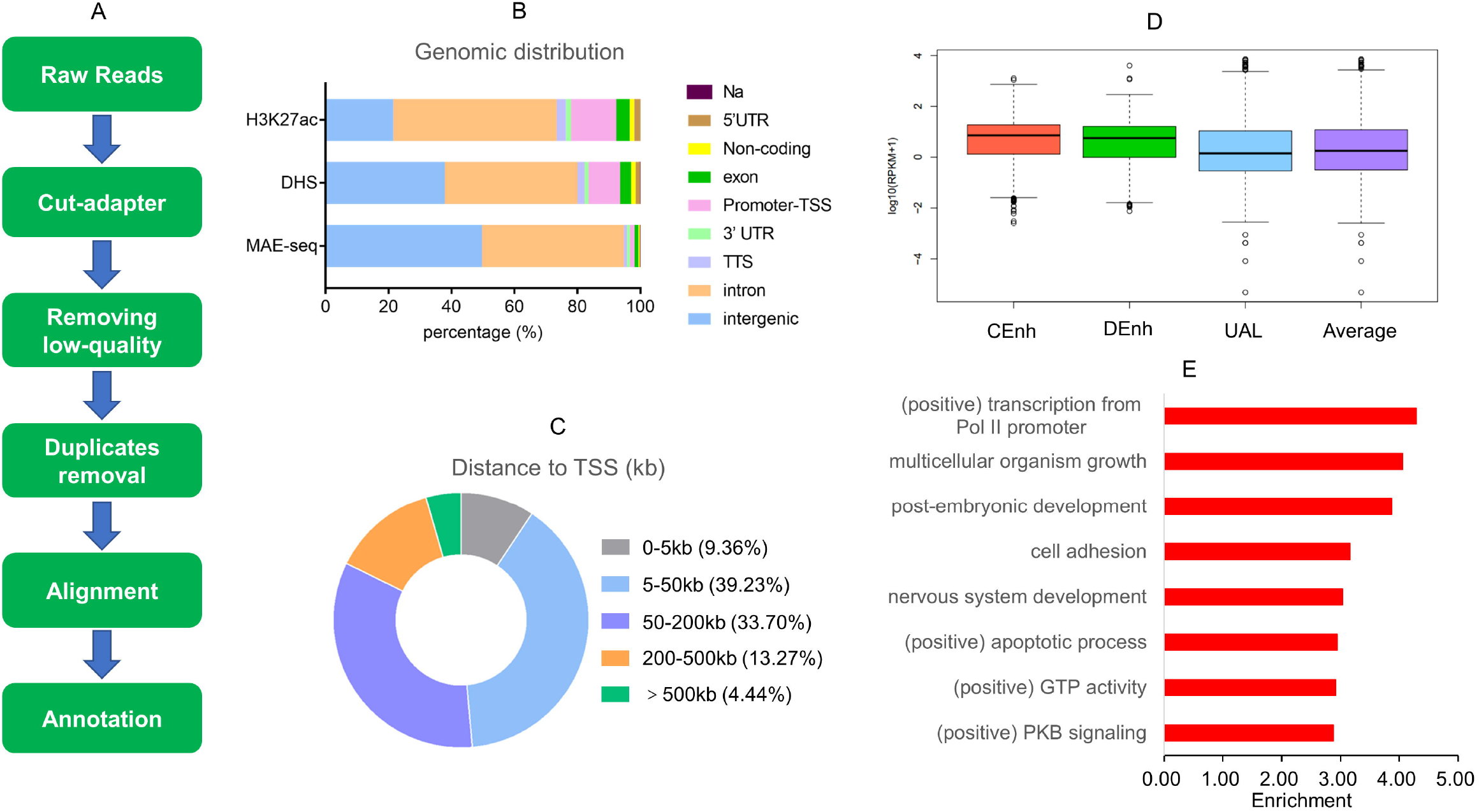
Genomic analysis and GO. A: Brief analysis procedure of MAE-seq data. B: Distribution of unique-mapping reads in genome. C: Distance of mapped signal away from TSS. D: Expression of enhancers neighbored genes referred from RNA-seq data derived from HEK293 cells. CEnh: conventional enhancer (5-50kb away from TSS), DEnh: distal enhancer (50-200kb away from TSS), UAL: unannotated loci, Average: mean expression level. E: GO analysis of genes annotated by conventional enhancers.

**Table 1.** Data filtering.

| Raw data | Cutadapter | fastq_quality_filter | Fastuniq |
| --- | --- | --- | --- |
| 7,412,991 | 7,412,979 | 7,406,623 | 532,017 |

**Table 2.** unique mapped reads without duplication, deletion and insertion.

| Unique mapped<br>reads | Rmdup-unique<br>mapped reads | Unique-site |
| --- | --- | --- |
| 22,529 | 12,412 | 9,771 |

To determine their effects on neighbored genes (27% to 47% enhancers are supposed increasing expression on them^18^.), a joint analysis with RNA-seq data of HEK293 cells is performed (Gene Expression Omnibus (GEO), GSM3432525) (table S2). Comparing with UAL (unannotated loci) and average level, genes neighboring the CEnh or DEnh locus have higher expression level (p<0.01) (Fig. 3D). It suggests that the active enhancer which identified by MAE-Seq play a positive role in transcription regulation.

GO analysis of target genes shows that many genes are enriched in the biological process of transcriptional activation (most enriched process which is also found in DNase-seq and ChIP-seq datasets) (Fig. S2) and some other positively regulated process (Fig. 3E) (Table S3).

### Cross-comparison and data validation

The enriched signals of MAE-seq annotated enhancers (9771) are analyzed in DNase-seq and H3K27ac ChIP-seq datasets derived from HEK293 cell (ENCODE, ENCSR000EJR, ENCSR000FCH) (Table S4 and S5). Loci of 618 (6.32%) enhancers are found to overlap with regions annotated by these datasets (Table 3). Moreover, parts of them locate within the range of 5kb regions enriched with DHS (DNase Hypersensitive Site) (26.66%) or H3K27ac (13.66%) signals (Fig. 4A and 4B) (Table 4). This implies that these enhancers are near regions predicted by ENCODE datasets. Moreover, distribution analysis result also indicates that genomic feature of MAE-seq enhancers is similar with which derived from previous methods (Fig. 3B).

**Figure 4.**
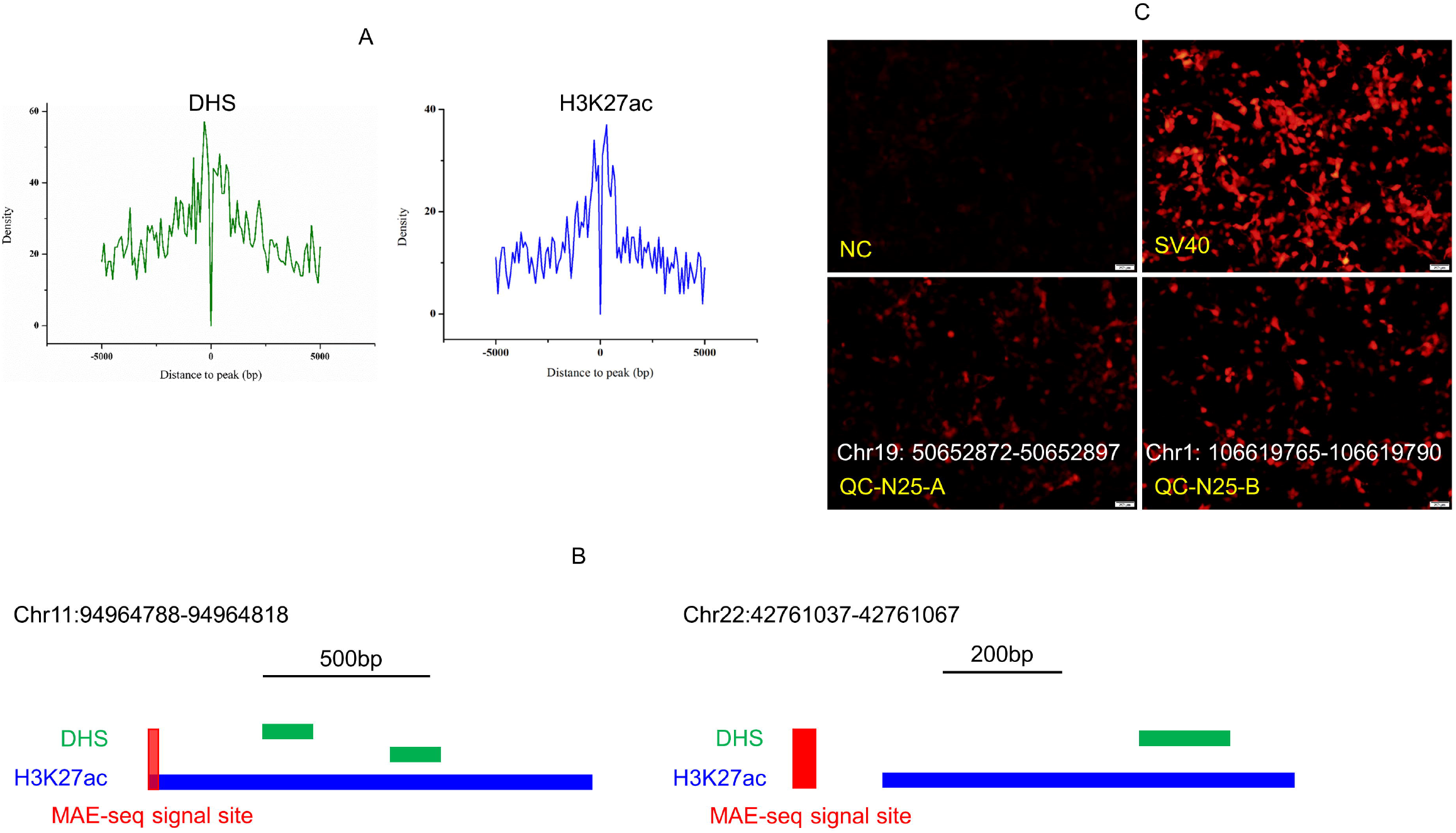
Epigenetic characteristics and validation of identified enhancers. A: The distribution curve of MAE-seq annotated enhancers which are 0-5kb away from DNase-seq peaks or H3K27ac ChIP-seq peaks. B: Example of MAE-seq annotated enhancer covered with or unmarked by H3K27ac ChIP-seq signal. C: Fluorescent signal of enhancers for validation. NC: cells transfected with empty vector, SV40: the positive control, QC-N25-A and QC-N25-B: two enhancers for validation. The length of the scale is 200 microns.

**Table 3.**
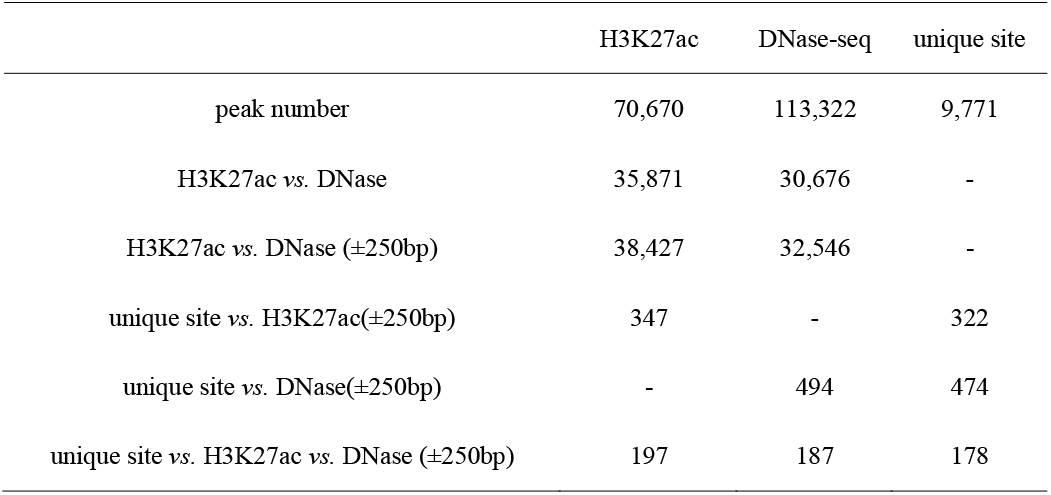
MAE-seq, DNase-seq, and chromatin mark enrichment.

**Table 4.** Separation distance of peaks between MAE-seq and DNase-seq or chromatin mark.

| distance to peak | 3kb | 5kb | 10kb | 20kb | 30kb | 40kb | 50kb | 100kb |
| --- | --- | --- | --- | --- | --- | --- | --- | --- |
| H3K27ac | 962 | 1,335 | 2,104 | 2,676 | 3,925 | 4,548 | 5,011 | 6,304 |
| DNase-seq | 1,828 | 2,605 | 3,986 | 5,617 | 6,573 | 7,165 | 7,593 | 8,586 |
| H3K27ac/total (%) | 9.85 | 13.66 | 21.53 | 27.39 | 40.17 | 46.55 | 51.28 | 64.52 |
| DNase-seq/total (%) | 18.71 | 26.66 | 40.79 | 57.49 | 67.27 | 73.33 | 77.71 | 87.87 |

Besides, there are 9153 (93.68%) enhancers have no overlap with DNase-seq and H3K27ac ChIP-seq datasets (Table 3). To validate the reliability of these loci, we select ten sequences derived from annotated loci randomly, and detect their activation in cells (Fig. 4C) (Table S6). The positive rate is more than 60% (Fig. S3) which suggests that this method can effectively annotate enhancers in genome. In addition, it also implies that large amounts of UREs^9^ without epigenetic features tend to be spotted by MAE-seq (Fig. S3).

## Discussion

Existing methods such as ChIP-seq, DNase-seq and STARR-seq, only enable rough enhancer predictions^19^. With these data, it is difficult to precisely define enhancer sequences less than 147bp, due to the structural characteristics of nucleosome. To avoid the limitation, MAE-seq is developed in this study based on short random DNA libraries and fluorescence-assisted screening. Thousands of active enhancers are archived in 25 bp length, which could dramatically improve the annotation details of regulatory elements revealed by ENCODE project. Moreover, large amounts of UREs have also been annotated across the genome. It implies more efforts might still be needed to comprehensively annotate the genome.

With MAE-seq datasets, we can deconstruct different regulatory elements, such as super enhancer. With the known definition, super-enhancers usually spread tens of kilobytes (kb) with signal clusters (Fig. S4)^14^, which might be complicated to study its molecular mechanism based on these^15^. With our data, it has the potential to expose the enhancer’s distribution within the region. In the long run, this would be helpful to clarify their definition and explore their mechanism on cell identity and etc^20–23^. Besides that, MAE-Seq identified enhancers provide an effective tool for functional studies. With current other methods, enhancer sequences are ranged from ~200bp to a few kb. Considering the limited carrying capacity, it is a bit challenging to clone and analyze the function of them with available vectors, especially for the multiple enhancers which have the same target gene. Traditionally, multiple signals must be validated in segments^24^. It is inefficient and time-consuming. With MAE-Seq identified enhancers, multiple enhancers in tens bp could be analyzed in the same time. This might facilitate the related gene functional studies.

In this study, we have test the MAE-seq method with different lengths. It remains as challenge for their spatiotemporal studies. We will further optimize the procedure and compare the datasets across species and stages. Hopefully, we could decipher the accurate enhancer sequences across the genome in the future. At the same time, we are actively testing and trying to apply this strategy for other regulatory element categories experimentally, such as transcription factor binding sites^25,26^. Besides that, with the identified UREs, we are wondering its function model behind it because of their lacking epigenetic features, especially in three-dimensional pattern.

In conclusion, MAE-seq provides a *de novo* way to further annotate the genome. At the same time, it might open a gate for enhancer mediated gene functional studies. We hope it could advance our knowledge on the genome with those findings.

## Methods

### Construction of reporter vector

The reporter vector, pMX-mP-mCherry based on the modified pMX-GFP retroviral vector (Cell Biolabs, USA), is used in the study. Mini-promoter and mCherry (miniP-mCherry) fragments are obtained from modified pGL4.23 luciferase reporter vector (Promega, USA). Briefly, PGL4.23 is digested with XbaI/NcoI restriction enzymes to remove the original coding sequences of luc2 reporter gene and replace it with mCherry. The miniP-mCherry fragment is then amplified with primers containing NcoI or NotI restriction enzymes. Synthetic DNA fragment is cloned into NcoI and NotI digested pMX-GFP vector. The resulting screening vector is named pmx-mchery.

### Generation of random DNA library

Random DNA library is constructed following a two-step process (DNA fragments synthesis and Seamless Assembly Cloning) (Figure 1B). DNA consisting a series of different lengths DNA fragments and of 30bp homologous arm at its two ends is synthesized by GENEWIZ (Suzhou, China), Synthesized random DNA is then inserted into reporter vector through homologous recombination methods following Clone Express J One Step Cloning Kit protocol (Vazyme, Nanjing, China).

Sanger sequencing is conducted to check the quality of the library clone. A total of ten single clones of E. coli are randomly selected from luria broth (LB) agar plate. Each single clone is then transferred to LB liquid and incubated at 37°C for 12-16 hours in a shaker incubator. The incubated bacterial suspensions are sent to company (Sangon Biotech, Shanghai, China) for sequencing. Bacterial suspension of library clones with high quality sequence is further used for plasmid extraction.

### Cell culture and transfection

Cell cultures are inoculated into 100 mm DMEM (ThermoFisher; Cat. No. 10566024) containing 10% FBS (Gibco; Cat. No. 10437028) and 1% Penicillin Streptomycin P/S (Sigma; Cat. No. P0781) at 37 □ cell incubator. Transfection of plasmid libraries (4μg DNA/1×10^6^ cells) is performed with cells at 70-80% confluence using HD Transfection Reagent (FuGENE; Cat. No. E2311). Fluorescence cells are then collected 24 hours after transfection.

### Establishment of a screening library for active enhancers

DNA is extracted from the collected fluorescent cells and amplified for 7 cycles using primers containing reporter vector sequence (forward: TGCAGGTGCCAGAACATTTC, reverse: GTGGCTTTACCAACAGTACC). The amplified products are electrophoresed in 2% agarose gel for 35 min (120v,130mA) and recovered through gel recycling. The recovered product is subjected to 4 cycles of amplification using KAPA kit (KAPA, Cat. No. KK2620) and sent for high-throughput sequencing.

### Identification of active enhancers

The raw data (raw reads) of fastq format are firstly trimmed by cutadapt (v1.16) to remove adapter and homologous arm sequences ^27^. Then, low quality reads are filtered by fastx-toolkit (http://hannonlab.cshl.edu/fastx_too1kit/)^28^. Only the sequences which the number of bases with a minimum Phred-score quality score of 20 accounted for more than 50% of the total would be saved. Next, the sequences are processed through Fastuniq (https://sourceforge.net/projects/fastuniq/) to remove the duplicated sequences. Finally, the remaining reads which have the same length as that we inserted are picked out and aligned to hg19 reference genome using Bowtie 2 version 2.1.0 with parameters -D 20 -R 3 -N 0 -L 17 -i S,1,0.50 --end-to-end. The unique mapping sequences which have no deletion and insertion compared with the reference are kept^29^.

### Enhancer distribution in the genome

The unique mapping regions in the genome are annotated by ChIPseek^30^. To study the enhancer distribution in genome, the genome is annotated in the following categories: promoter-TSS, TTS, intergenic, exon, intron, 5’UTR, 3’UTR and non-coding. Each enhancer is sorted into different category if its summit falls within sequence of a specific category. At the same time, the distance between enhancer and proximal TSS is computed.

### MAE-seq, DNase-seq, and chromatin mark enrichment

DHS and histone modification data, including H3K27ac are retrieved from ENCODE. To study the associations between the unique mapping regions and DNase-seq, H3K27ac peaks, we extend the region of peak by ± 250bp, and use bedtools to calculate the proportion of overlap between the unique mapping regions and the extended peaks^31^. The distribution of MAE-seq enhancers around the center of peak from ChIP-seq(H3k27ac) and DNase-seq is computed and displayed by R.

### Statistical analysis

We use R for all statistical analysis.

## Supplementary Data

Supplementary Data are available online.

## List of abbreviations

ATAC-seq: (Assay for Transposase-Accessible Chromatin using sequencing),
(bp): base pairs,
ChIP: (chromatin immunoprecipitation),
(CEnh): conventional enhancer,
(DEnh): distal enhancer,
DHS: (DNase Hypersensitive Site),
DNase-seq: (DNase I hypersensitive sites sequencing),
ENCODE: (Encyclopedia of DNA elements),
FAIRE-seq: (formaldehyde-assisted isolation of regulatory elements sequencing),
FI: (fluorescence intensity),
GEO: (Gene Expression Omnibus),
HTS: (high-throughput sequencing),
MAE-Seq: (massive active enhancer sequencing),
STARR-seq: (self-transcribing active regulatory region sequencing).

## Availability of data and material

Sequence data are submitted to the GEO repository (GSE129453). DNase-seq, H3K27ac data and RNA-seq data are achieved from ENCODE (ENCSR000EJR, ENCSR000FCH) and GEO (GSM3432525) dataset respectively.

## Authors’ contributions

Y.Z. and L.H. conduct and design the experiments. X.Z. performs the experiments. Q.L. conducts the bioinformatics work. X.Z. and Q.L. participates in discussions and give good advice. The manuscript is written by L.H. and Y.Z. All authors read and approve the final manuscript.

