## Supplemental file for "Start from Scratch: Precisely Identify Massive Active Enhancers by Sequencing"

The supplementary file contains files as follows:

Figure S1. Construction of sequencing library.

Figure S2. GO analysis of highly expressed genes activated by conventional enhancers.

Figure S3. Data validation.

Figure S4. ChIP-Seq binding profiles for Med1 at a super-enhancer locus in mouse embryo stem cell.

Table S1. N25 unique site annotation.

Table S2. Relationship between unique site and gene expression.

Table S3. GO analysis of MAE-seq, DNase-seq and H3K27ac enhancers.

Table S4. H3K27ac overlap with MAE-seq.

Table S5. DNase-seq overlap with MAE-seq.

Table S6. Sites information for data validation.

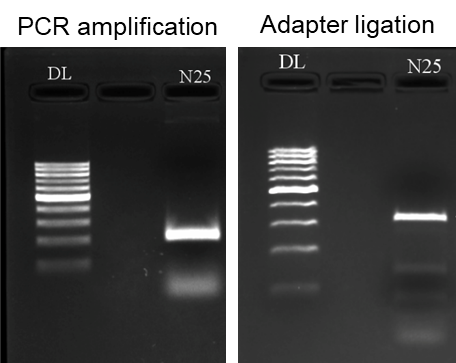

Figure S1. Construction of sequencing library.

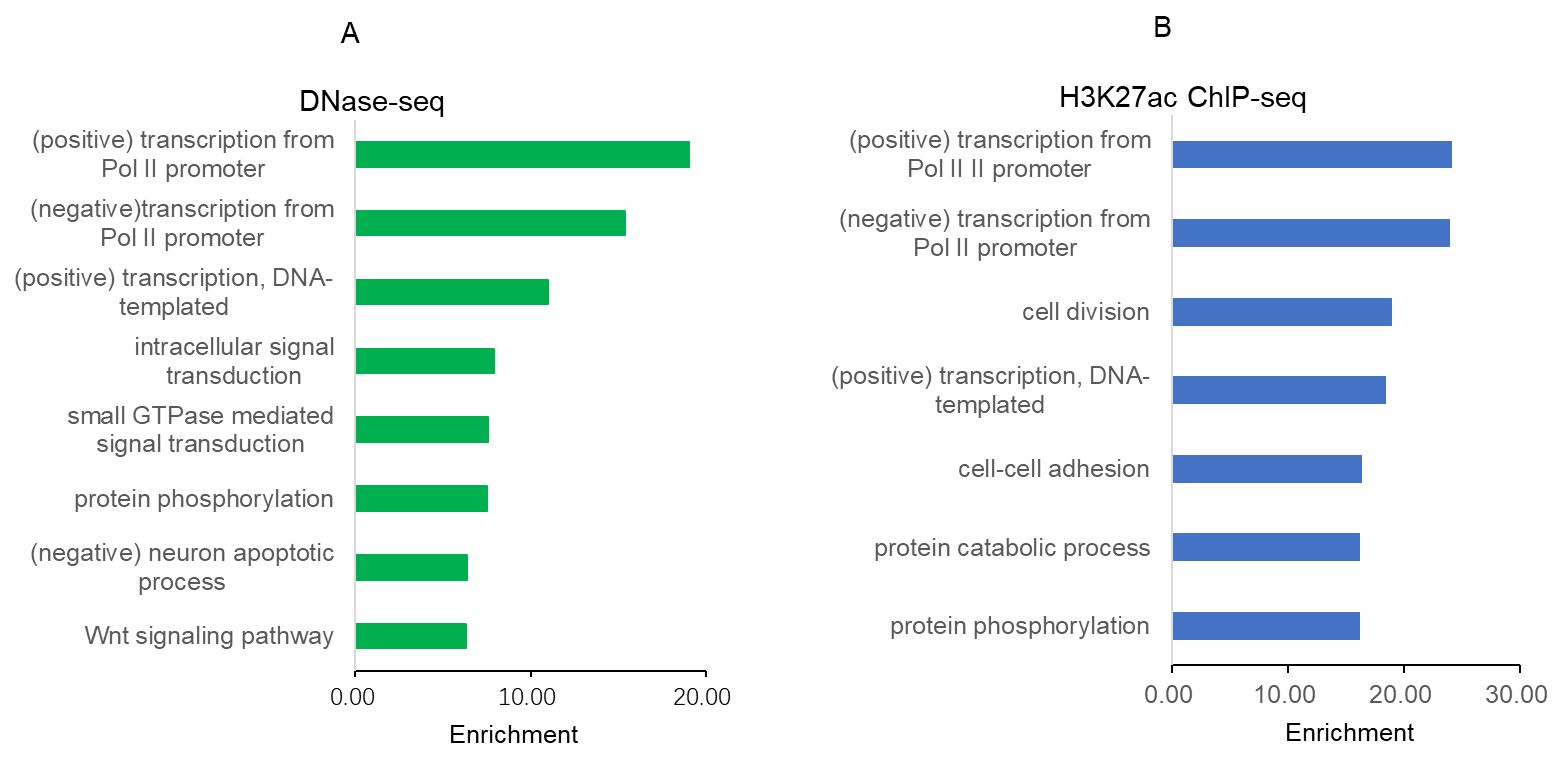

Figure S2. GO analysis of highly expressed genes activated by conventional enhancers.

GO analysis of genes related to enhancers predicted by DNase-seq and ChIP-seq.

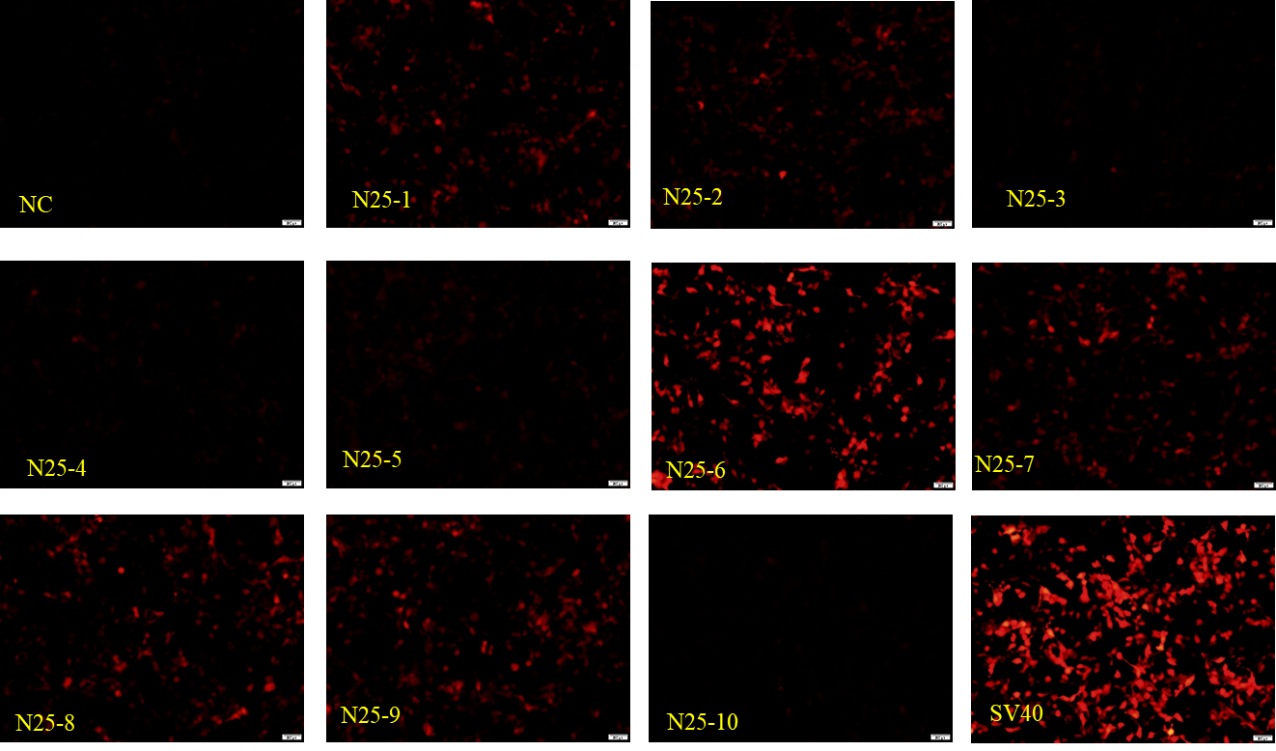

Figure S3. Data validation.

Data validation corresponds to Fig. 4 in the main text.

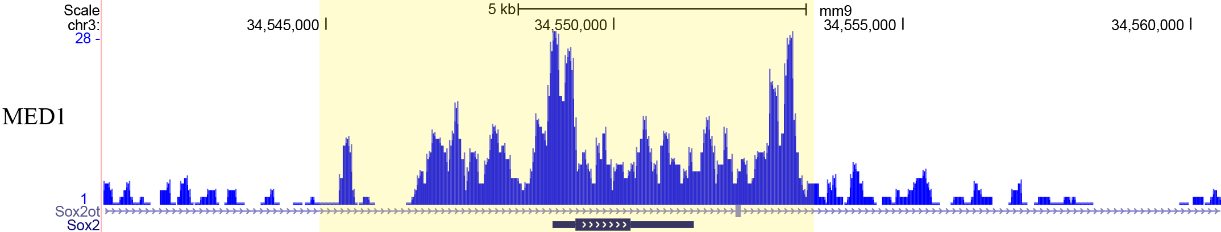

Figure S4. ChIP-Seq binding profiles for Med1 at a super-enhancer locus in mouse embryo stem cell.
